# Network rewiring promotes cooperation in an aspirational learning model

**DOI:** 10.1101/2021.09.16.460462

**Authors:** Anuran Pal, Supratim Sengupta

**Affiliations:** Department of Physical Sciences, Indian Institute of Science Education and Research Kolkata, Mohanpur Campus, Mohanpur-741246, India

## Abstract

We analyze a cooperative decision-making model that is based on individual aspiration levels using the framework of a public goods game in static and dynamic networks. Sensitivity to differences in payoff and dynamic aspiration levels modulate individual satisfaction and affects subsequent behavior. The collective outcome of such strategy changes depends on the efficiency with which aspiration levels are updated. Below a threshold learning efficiency, cooperators dominate despite short-term fluctuations in strategy fractions. Categorizing players based on their satisfaction level and the resulting strategy reveal periodic cycling between the different categories. We explain the distinct dynamics in the two phases in terms of differences in the dominant cyclic transitions between different categories of cooperators and defectors. Allowing even a small fraction of nodes to restructure their connections can promote cooperation across almost the entire range of values of learning efficiency. Our work reinforces the usefulness of an internal criterion for strategy updates, together with network restructuring, in ensuring the dominance of altruistic strategies over long time-scales.

Maintaining a public resource requires sustained cooperation through contributions by community members who benefit from it. Yet, a selfish individual who refuses to contribute can enjoy the benefits without paying the cost of sustaining the public good. If however, too many members of the community act selfishly, the public resource collapses to the detriment of all. The public goods game highlights such a social dilemma and provides a framework for exploring different mechanisms of strategic decision-making that allow cooperation and consequently the public good to be sustained. Among many mechanisms, the reorganization of social ties has been shown to be effective in promoting cooperation in PGG. However, the efficacy of most mechanisms in sustaining cooperation rely on individuals updating their strategy on the basis of information about the contributions of other members of the community. Often such information is either not forthcoming or cannot be effectively utilized. An alternative low-information model of behavioral updating relies on a comparison between the actual benefit received and the benefit aspired for. Individuals tend to retain their strategy if they are satisfied with the benefit received and change their strategy if they are unsatisfied. We show that such a simple reinforcement learning model along with modest restructuring of social ties over time can allow cooperation to be sustained. Our work shows that a low-information strategy-update model can be very effective in ensuring dominance of cooperators in social dilemmas.

## I. INTRODUCTION

Evolutionary game theory provides a useful framework for understanding how cooperation can be sustained in a population where individual interests are at cross purposes with that of the group. A classic example of such a social dilemma is often presented through the example of a public goods game (PGG) where individual exploitation of a public good at the expense of other members of the community can ensure short-term gains for the selfish individual but eventually lead to the decimation of the public good, a consequence that is detrimental to all members of the community^1^. It is therefore imperative to uncover either behavioral or policy-based mechanisms that can avoid socially costly outcomes such as the “tragedy of the commons” in the long run.

Many of the mechanisms^2,3^that have been proposed for explaining the sustenance of cooperators are reactive in the sense that they specify an appropriate response to the altruistic or selfish actions of others. The response itself can change through individual learning that can be conditioned on a variety of factors like the latest payoff, strategy and wealth environment^4–7^, social norms^8–15^and even personal values. However, most evolutionary game-theoretic models emphasize the importance of latest-payoff in shaping the evolution of individual decisions which typically occurs through the comparison of the focal player’s payoff with that of a randomly chosen neighbor^16,17^. Many strategy update rules also rely on information about last-round strategies of other members of the population. Such information may not always be forthcoming. Even if they are available, their usefulness will be limited by the information processing capabilities of individuals especially in large populations. It is therefore natural to explore the consequences of low-information decision heuristics on the evolution of cooperation in social dilemmas. This has motivated the study of alternative, low-information strategy update mechanisms such as aspiration dynamics^18–21^where individuals change their strategy on the basis of a comparison between their payoffs and personal aspiration level. Cooperation levels in social dilemmas such as a PGG is also influenced by the structure of the underlying social network as well as evolution of the underlying network structure due to the reorganization of social ties. The coevolution of individual decisions as well as social ties^22–31^can often favour the proliferation of cooperation in a manner that depends on the details of the network restructuring process.

Reinforcement learning models have been extensively studied^32–38^in the context of repeated games, involving just 2 persons, to show how cooperation can be sustained even though such a strategy is not a Nash equilibrium^35,36,39^. In such models, called the Bush-Mosteller^40^ (BM) reinforcement learning models^36^, an action is likely to be repeated if its outcome (manifest through received payoff) ensures the satisfaction of the player. Satisfaction is determined by comparing the payoff with the aspiration level. The effect of learning is incorporated through a changing aspiration level^35,36,39,41,42^with learning efficiency parametrized by a habituation parameter (see Methods section below). Even though, most of analysis of Macy and Flache^36^were carried out for 2-person repeated games with fixed aspiration level, they found that an adaptive aspiration level characterized by a single (low) value of the habituation factor is counterproductive for sustaining mutual cooperation. Masuda and Nakamura^41^ modified the BM model of Macy and Flache to make the satisfaction sensitive to the difference between the actual and aspired payoff. They found that mutual cooperation can be sustained for even moderate values of learning efficiency and such BM players can even outcompete many other memory-one strategies. Roca and Helbing^43^ proposed a different type of low-information, learning model where the aspiration level changes between the minimum and maximum payoff in a manner that depends on the individual’s greediness. Their results, recently extended to evolving complex networks^44^, showed that cooperation and social cohesion (measured by the fraction of individuals who do not change either strategy or social neighborhood relative to the last time-step) can be sustained at moderate levels of greediness even in the absence of information about strategies of connected neighbours.

Despite progress in our understanding of the impact of reinforcement learning on evolution of cooperation, many questions still remain unanswered. How is the extent of cooperation affected by learning efficiency as well as the underlying network topology in social dilemmas like PGG’s? To what extent does reorganization of the underlying network through change in social ties affect cooperation levels? Is it possible to identify the microscopic mechanisms that allow for dominance of altruistic strategies? In this paper, we adopt the model proposed in^41^and use the framework of a PGG in both static and dynamic networks to investigate the consequences of reinforcement learning on the proliferation of altruistic strategies. We find that cooperation can not only be sustained in the population but also dominate over selfish strategies as long as learning efficiency is not too high in static network-structured populations. This is achieved *despite* large short timescale fluctuations in cooperator fractions. Our analysis of short time-scale dynamics reveals that the cooperator-dominated and defector-dominated phases are characterized by distinct types of cyclic transition dynamics that occurs between different categories of cooperators and defectors. Allowing network restructuring can lead to dominance of cooperators for even higher learning efficiencies compared to the threshold obtained for static networks.

## II. METHODS

The model consists of a network of *N* nodes. Each node represents an agent (*i*) who plays (*k*_*i*_ + 1) modified PGG with all her *k*_*i*_ neighbors, once as a focal player and the remaining *k*_*i*_ times as a neighbor of a focal player, in each round. If an agent decides to cooperate as a focal player, she pledges a fixed amount of money (*m* = 1) to each connected neighbor, which after getting multiplied by a synergy factor *r*, is distributed to all her neighbours, regardless of their actions. A selfish agent does not donate any money to her neighbors. Thus, the payoff of an individual player is:

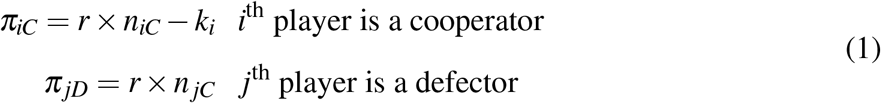

where, *n*_*iC*_ denotes the total number of cooperator neighbors and *k*_*i*_ denotes the degree of the *i*^th^ player.

### A. Decision making

An agent’s propensity to cooperate at time (*t* + 1) depends on her satisfaction level (*s*_*i*_(*t*)) after the last round. The latter depends on the relative difference between the latest payoff (*π*_*i*_(*t*)) and aspired payoff (*A*_*i*_(*t*)).

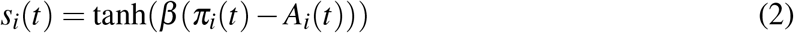

*β* captures the sensitivity of the agents to the difference between received and the aspired payoff. The tanh function ensures that the satisfaction is bounded between, −1 ≤ *s*_*i*_(*t*) ≤ 1.

The satisfaction modifies the probability of cooperation in such a way t hat i f the l ast action (*C* or *D*) of the agent resulted in her satisfaction *s*_*i*_(*t*) *>* 0, the propensity of continuing with that action is further reinforced in the next round^41^ as follows

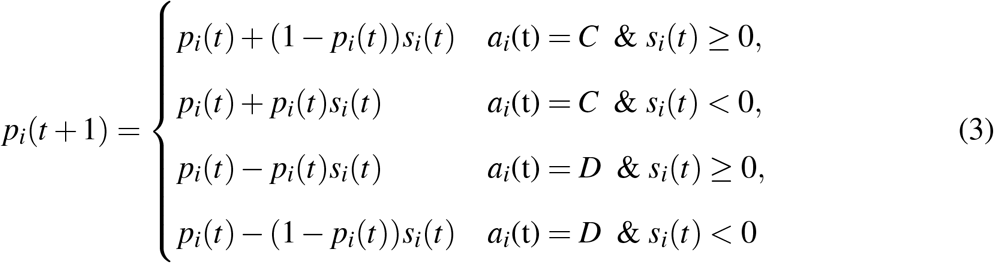

where *p*_*i*_(*t* + 1) gives the probability of cooperation of agent *i* in round *t* + 1 and *a*_*i*_(*t*) is the strategy employed by the *i*^th^ agent at time *t*. An implementation error (*ε* = 0.02) is considered to account for the trembling hand effect. The action *a*_*i*_(t+1) determined on the basis of probability estimated at time (t+1) is flipped with a probability *ε*. This prevents the system from getting trapped in suboptimal minima when the probability of cooperation is either 0 or 1.

The *i*^th^ agent updates her aspiration level in the next round in a manner that is determined by the extent to which she can learn from her past experiences. A perfect learner always updates her aspiration level to match her latest payoff, while an agent who is incapable of learning continues to retain her original aspiration level fixed at the beginning (*t* = 0) of the game.

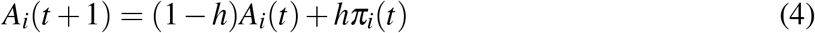

Learning efficiency is determined by the habituation factor *h*, which is an attribute of the population. Higher habituation values result in aspiration of players being primarily influenced by the payoff in the last round, whereas, lower habituation values ensure that the aspiration levels are significantly influenced by payoffs obtained in the near to distant past. At the beginning of every round, the aspiration level of every member is reset (for *h >* 0) on the basis of the pay-off received by her in the previous round according to Eq.4. This is followed by the PGG game that leads to new payoffs for every member. Subsequently, the satisfaction level of each member is determined by comparing her latest payoff with her aspiration level according to Eq.2. This satisfaction level of each member is used to estimate their probability of cooperation in the next round according to Eq.3. Cooperators and defectors are divided into two sub-categories each, that are distinguished by their individual satisfaction level. A satisfied/unsatisfied cooperator (SC/UC) is one that opts to cooperate in the next round after being satisfied/unsatisfied with her payoff (relative to her aspiration) obtained as a consequence of her action in the current round’s PGG. Similarly, a satisfied/unsatisfied defector (SD/UD) is one that opts to defect in the next round after being satisfied/unsatisfied with her payoff obtained as a consequence of her action in the current round. To better understand the collective dynamics, we also compute the average satisfaction of all those individuals who opt to cooperate in the next round and those who opt to defect in the next round after receiving payoffs on the basis of their actions in the current round (see Fig.1).

**FIG. 1.**
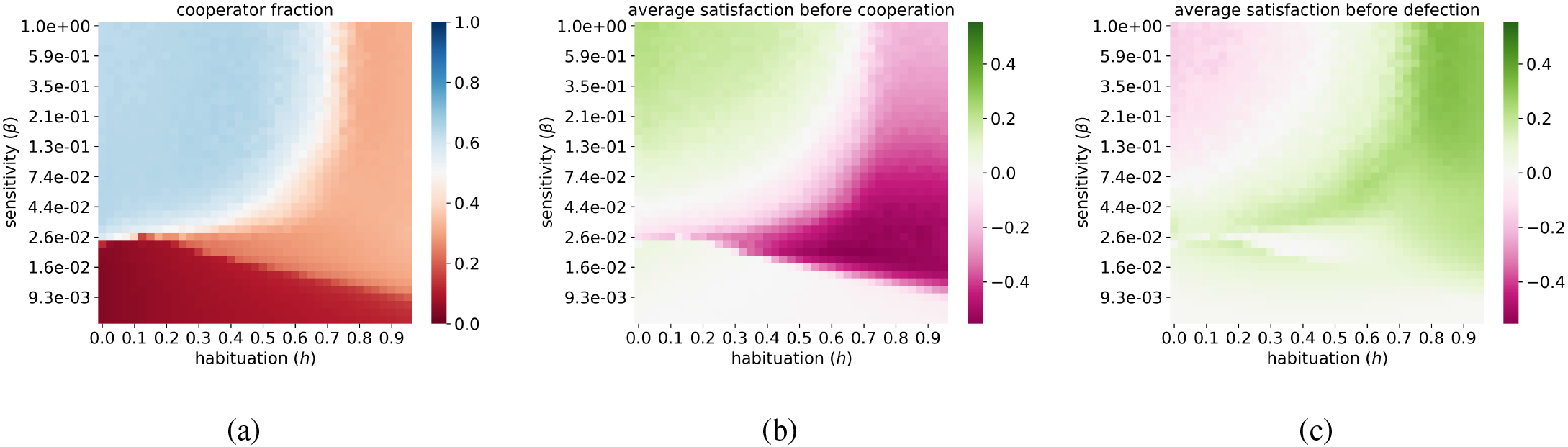
Heat map showing the (a) cooperator fraction; average satisfaction of all individuals who (b) cooperate in the subsequent round and (c) defect in the subsequent round; as a function of habituation (*h*) and sensitivity (*β*) for *N* = 500. Each pixel value is obtained by first averaging from round 250 to 750 for each trial and then averaging over 5 trials. The horizontal axis represents forty habituation values running from 0.025 to 0.975 and the vertical axis represents thirty-nine *β* values evenly spaced from 10^−2.2^ to 1 on a log scale. The other parameter values are *ε* = 0.02, and *r* = 2.

#### Initial conditions

The population consists of 500 agents constituting the nodes of an Erdős Re"nyi (ER) graph with 0.3 as the probability of a link formation between any two nodes. Initially, all players cooperate with a probability of 0.5. Consequently, approximately half of all the players defect whereas the other half cooperates, in the first round of the games. Because of the dynamic nature of the aspiration, the initial aspiration levels of the players have no effect on the dynamics after a few rounds, provided the habituation parameter is not vanishingly small. The initial aspiration levels of each player are set to the value *k*_*i*_ ×*m*× (*r*− 1)*/*2 where, *k*_*i*_ is the degree of the player and *m* = 1. This is the expected payoff of the *i*^th^ player with *k*_*i*_ number of connections, where all the players, including the focal player, cooperate with a probability of 0.5.

### B. Network rewiring

After every round of the game, each pair of players is given an opportunity to modify their connections with a probability *ρ*, i.e. approximately *ρ* fraction of the 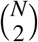 pair of players are allowed to either make a new connection if there wasn’t any or break an existing connection; before the next round of the game. We implement two different methods for restructuring social ties.

#### Rewiring based on strategy information

Rational players have a much higher tendency to connect with cooperators to increase their payoffs in the next round. Similarly, cooperators have a higher tendency to break links with selfish neighbors. Thus, allowing players to access information about their neighbors’ actions in the previous round induce individuals who have cooperated in the past-round to establish more connections than past defectors. The network rewiring model used in this section was inspired by behavioural experiments^7^ performed to analyse evolution of cooperation and wealth distribution using PGG. Each of the *ρ* fraction of the existing links are maintained or broken based on the action of one of the two participating players, chosen randomly^4,7^. If the partner of the decision-maker had defected in the previous round, the decision-maker, regardless of her own action in the previous round, chooses to break the connection with probability *p*_*b*_. On the other hand, if the partner had cooperated in the last round, the link is retained with probability *p*_*r*_. If the pair among the *ρ* fraction of pairs of players selected for rewiring had no link in the previous round, the action of *both* the players dictate whether a new link is established between them. A pair of previously unconnected cooperators have a high probability *p*_*m*_ of establishing a new link. A defector has a much lower probability of establishing a new link. If either or both players had defected in the last round, a new link is established between them with probability *p*_*e*_ and *p*_*s*_ respectively. Following Pathak et. al.^4^, we use *p*_*b*_ = 0.7, *p*_*r*_ = 0.87, *p*_*m*_ = 0.93, *p*_*e*_ = 0.3, *p*_*s*_ = 0.2.

#### Rewiring based on satisfaction

We also consider a form of network restructuring that does not require information about strategies of players. An individual can rewire her social ties based on her satisfaction. We have assumed that the willingness of players to make or retain a link is independent of the magnitude and depends only on the sign of a player’s satisfaction. A satisfied (*s*_*i*_ *>* 0) and an unsatisfied (*s*_*i*_ *<* 0) player decides to form a new link or retain an existing link with probability *p*_1_ and *p*_0_ respectively. Both players must opt to form a new (or retain an existing) link independently of the potential partner as well as past decisions. Hence, a pair of satisfied players (or unsatisfied players) form a new link or retain an existing one with probability 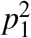 (or 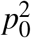). Similarly, a satisfied and an unsatisfied player make a new link or retain an existing one with probability *p*_1_ *p*_0_. We use *p*_1_ = 0.8 and *p*_0_ = 0.2. The results are robust to reasonable deviations from these values.

## III. RESULTS

### A. Static network

Fig.1(a) shows the phase diagram obtained by varying the habituation factor (*h*) and the sensitivity parameter (*β*). It reveals three distinct phases that are distinguished by the relative fraction of cooperators in the population. Cooperators dominate only when *β* is above a critical threshold (*β*_*c*_, whose value is dependent on the habituation factor) and the habituation factor below a critical threshold (*h*_*c*_). Very low values of *β*, when an agent’s satisfaction becomes almost independent of her dynamic aspiration, are characterized by complete dominance of defectors across all learning abilities determined by *h*. On the contrary, high sensitivity (large *β*) can amplify small differences between payoff and aspiration to drive satisfaction towards its saturation values of *±*1. This increases the likelihood of both cooperators and defectors retaining their previous strategies if they were satisfied or switching to the competing strategy if they were dissatisfied in the last round. For large *β* ; as learning efficiency (*h*) increases, the aspiration level of individual agents are predominantly determined by their latest payoff, defectors again start dominating the population. This behavior can be better understood by further distinguishing between sub-groups of satisfied and unsatisfied cooperators and defectors. As is evident from Fig.1(b)-(c), the dominance of cooperators or defectors is dependent on their satisfaction levels. For large *β*, as long as learning efficiency is below a critical threshold, cooperators dominate the population because they are satisfied on the average. As *h* increases, fluctuating aspiration levels that are now primarily dependent on latest payoff results in most cooperators becoming unsatisfied, leading to a dynamic equilibrium phase that is dominated by mostly satisfied defectors. For marginally low values of beta, individual satisfaction depends not only on the sign but also on the magnitude of the difference between the aspiration and payoff. When that difference between aspiration and payoff is small, the individual satisfaction can be close to 0. If that difference is significant, even at critical beta, satisfaction can take on intermediate values resulting in the average satisfaction values for defectors averaging out to zero (Fig.1c) due to the roughly similar levels of abundance of both UD and SD players.

Fig.2a shows how the different categories of cooperators and defectors change with *h*. SC dominate the population as long as *h* is below a certain threshold (*h*_*c*_ *≈* 0.7); beyond which, SD starts dominating. The underlying strategy dynamics responsible for the phase diagram shown in Fig.1 can be understood in terms of the transitions between the different categories of cooperators and defectors and the manner in which learning ability (parametrized by *h*) affects strategy shifts in individual agents by determining how an agent responds to differences between her latest and aspired payoff.

**FIG. 2.**
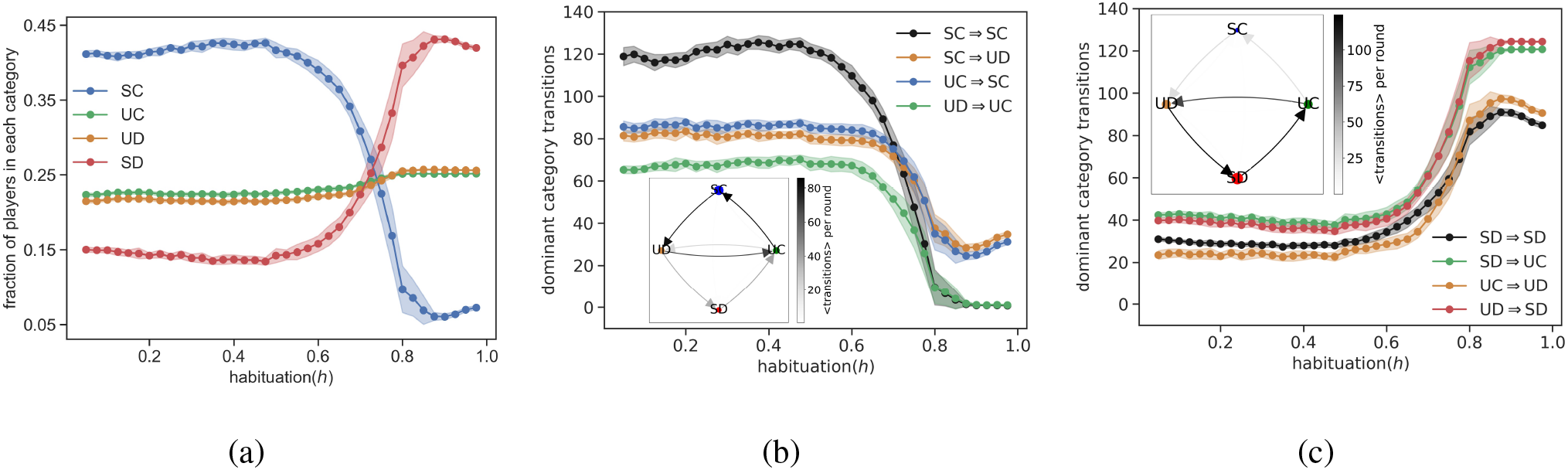
(a) Change in fraction of players in each category with habituation (*h*). (b) Dominant category transitions at low values of habituation. (c) Dominant category transitions at high values of habituation. All the simulations are time averaged from round 500 to 1000 over 25 trials. The shaded region denotes one sigma deviation from the mean. The figures in the inset of (b) and (c) denote the distinct transition cycles in the moderate (*h* = 0.4) and high (*h* = 0.9) habituation regimes respectively. The dominant transitions are denoted by darker shades of arrows and the fraction of the population in the different categories are denoted by the size of the circles. The other parameter values are *N* = 500, *β* = 1, *ε* = 0.02, and *r* = 2.

The aspiration for round *t* + 1 can be expressed in terms of the initial aspiration and the entire payoff history of the player.

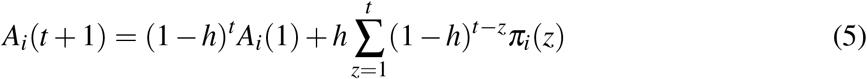

Low to moderate values of *h* ensures considerable contribution from payoffs obtained prior to the last round’s payoff, *A*_*i*_(*t* + 1) ≈ *hπ*_*i*_(*t*) + *h*(1 *−h*)*π*_*i*_(*t* − 1) + *h*(1 −*h*)^2^*π*_*i*_(*t −* 2). This leads to slower modulation of individual aspiration compared to the high learning efficiency scenario and affects transitions between different categories of cooperators and defectors. Fig.2b,c shows the dominant transitions between categories (i.e. the strategy shifts whose numbers change significantly) as *h* increases from a low to high value. For the *h* < *h*_*c*_ region, when payoffs received in the near to distant past determine current aspiration levels, the transitions that dominate form a cycle (SC→UD→UC→SC) as evident from Fig.2b. This suggests that some individuals who had cooperated after being satisfied with their payoff in the previous round become unsatisfied with their latest payoff and turn to defection. However, they quickly realize that employing such selfish strategies leaves them unsatisfied relative to their aspired payoff and eventually revert to cooperating, following which the cycle is repeated. Apart from such cyclic transitions, the other dominant process (SC→SC) implies that a large fraction of cooperators retains their strategy because they are satisfied with their payoffs as indicated in Fig.1b.

The situation changes dramatically for (*h* > *h*_*c*_) when satisfied defectors tend to retain their strategies as manifest through high SD→ SD transitions. Moreover, even if some defectors switch to an altruistic strategy after being unsatisfied with their latest payoff, their dissatisfaction level in this state soon makes them change back to a selfish strategy as characterized by the cycle SD→ UC→ UD→ SD (see Fig.2c).

The equilibrium discussed in previous paragraph is a dynamic one obtained by time-averaging over several rounds. Even for *h* < *h*_*c*_, the transitions between categories ensures that the cooperator fraction fluctuates dramatically over short time-scales. In fact, the category transition dynamics is better understood by analysing the short time-scale behavior of the individuals belonging to different categories as well as the total aspiration levels.

Fig.3a shows the overall cooperator fraction as well as the dominant transition cycle (SC→UD→UC→SC) in the *h* < *h*_*c*_ regime for a short period of time. The fall in cooperator fraction is preceded by increase in SC→UD transitions. This transition is triggered when the increasing aspiration of cooperators exceed the payoffs, which decrease due to implementation error (top panel of Fig.3(b)). However, the large number of resulting defectors are mostly unsatisfied since their payoffs fall below their aspiration levels (bottom panel of Fig.3b), leading to a peak in the UD→UC transitions that follows the peak in the SC→UD transitions (see Fig.3a). The cooperators find that their altruistic behavior increases their payoff beyond their aspirations, leading to a peak in the UC→SC transition shown in dark blue in Fig.3a. This marks the beginning of the short period of dominance of cooperators in the population before the cycle is repeated. In this regime, the relatively slow change in aspiration levels, allow a large fraction of cooperators to persist for longer periods of time ensuring a high value for the time averaged fraction of cooperators.

**FIG. 3.**
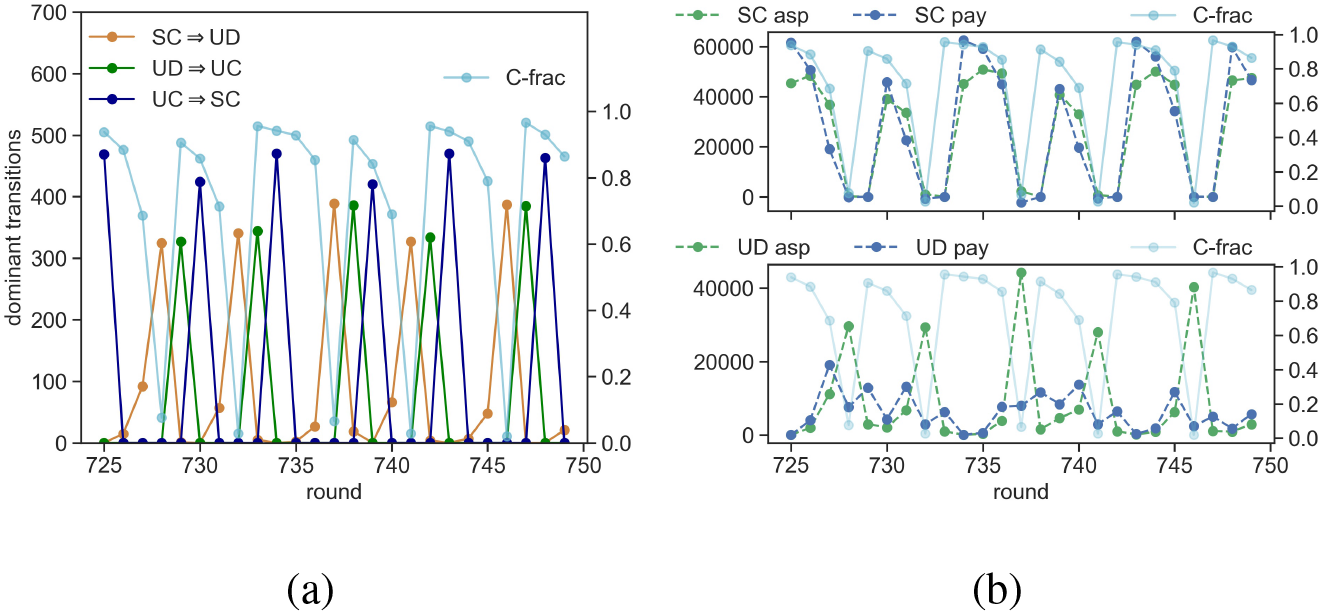
Short time-scale evolution of the (a) the cooperator fraction (C-frac) and the dominant transitions; total aspiration and total payoff of individuals beloging to the SC and UD categories for *h* = 0.4. The other parameter values are *N* = 500, *β* = 1, *ε* = 0.02, and *r* = 2.

A different type of cyclical dynamics is observed for *h* > *h*_*c*_, which is characterized by the pre-dominance of defectors. In that regime, the dominant transition cycles are (SD→UC→UD→SD) as shown in Fig.4a apart from the SD→SD transitions (see Fig.2c). While large fluctuations are seen in the cooperator fraction (Fig.4a), its time-averaged value remains quite low. An increase in cooperator fraction is strongly correlated with the increase in number of SD→UC transitions that are triggered because the payoff of defectors lag behind their aspirations (top panel of Fig.4b). But the altruistic act leaves the cooperators unsatisfied with their payoffs which lag behind their aspirations making the UC state unstable to further strategic transitions to defectors (bottom panel of Fig.4c). Changing to a selfish strategy result in large payoffs that exceed their individual aspiration levels and leads to the last step (UD→SD) in the transition cycle after which the cycle is repeated.

**FIG. 4.**
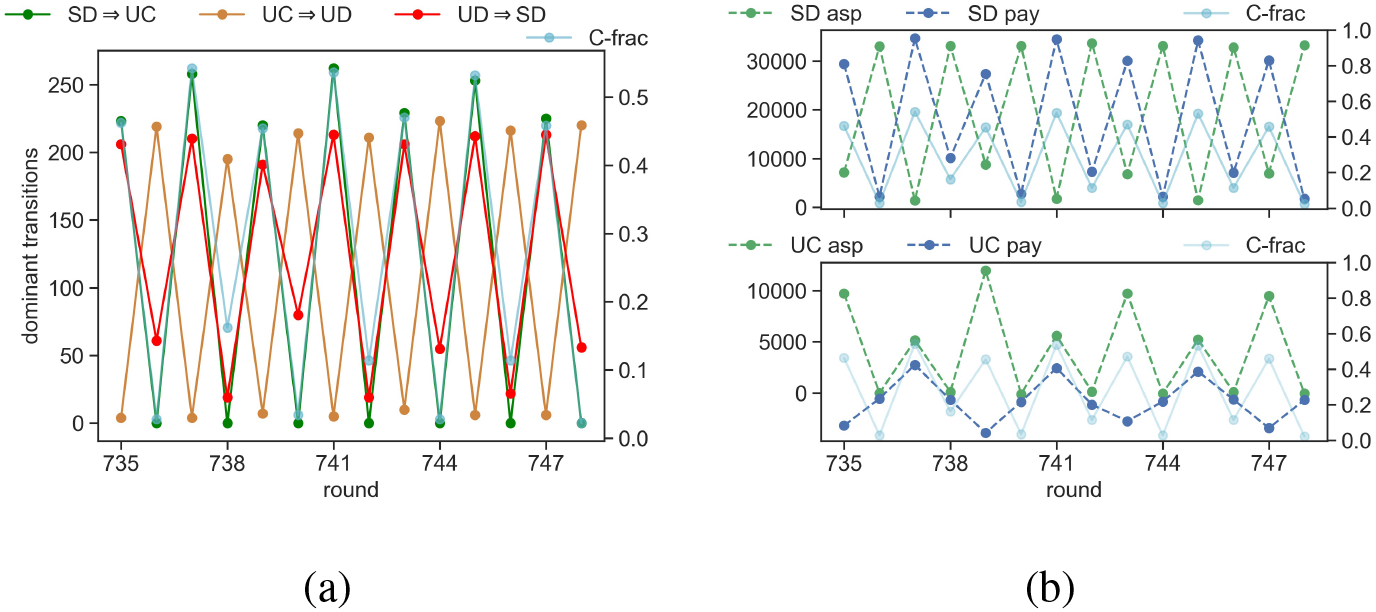
Short time-scale evolution of the (a) dominant transitions, cooperator fraction and aspiration & payoff of (b) SD & UC for *h* = 0.9. The other parameter values are *N* = 500, *β* = 1, *ε* = 0.02, and *r* = 2. C-frac denotes the fraction of cooperators.

### B. Dynamic network: Rewiring based on strategy information

The ability of agents to modify their social network by creating new and severing existing ties with other agents, on the basis of their knowledge of the last-round strategies of agents, allow cooperators to dominate the population even for high values of habituation parameter as can be seen in Fig.5a. The dominance of cooperators is typically correlated with their positive average satisfaction levels. Low to moderately high values of rewiring probability allow satisfied cooperators to thrive.

**FIG. 5.**
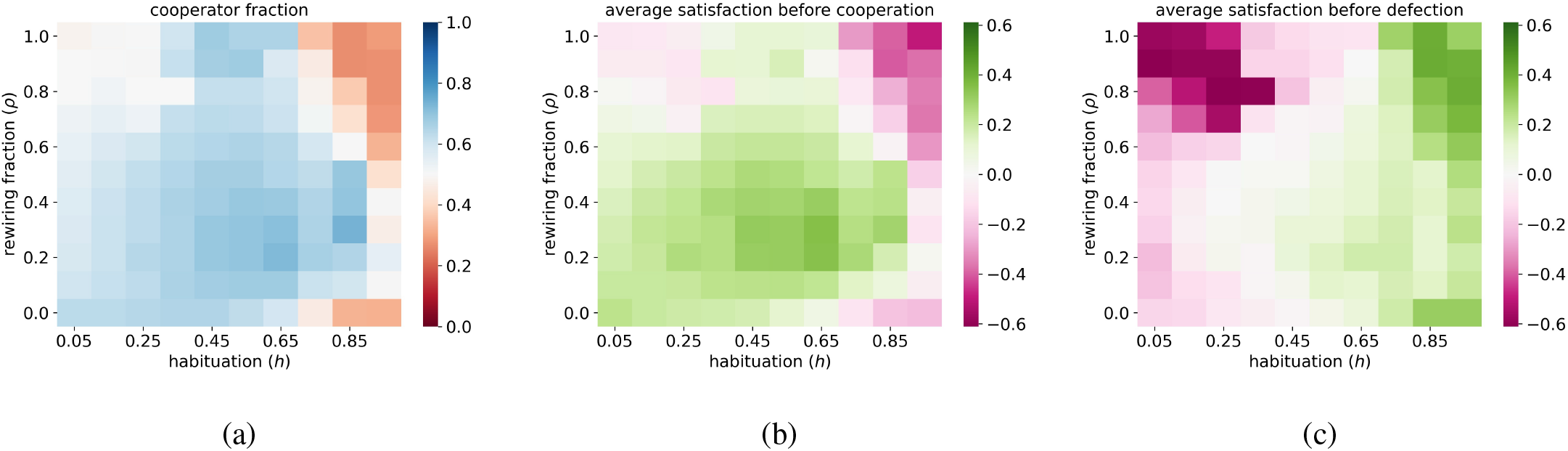
Heat map showing the (a) cooperator fraction; average satisfaction of all individuals who (b) cooperate and (c) defect in the subsequent round, as a function of habituation (*h*) and rewiring probability (*ρ*). The other parameter values are *N* = 500, *β* = 1, *ε* = 0.02, *r* = 2, *rho* = 0.3, *p*_*r*_ = 0.87, *p*_*b*_ = 0.7, *p*_*m*_ = 0.93, *p*_*e*_ = 0.3 and *p*_*s*_ = 0.2.

For moderate values of rewiring probability (0.1 ≤ *ρ* ≲ 0.7), a large fraction of cooperators is satisfied and therefore retain their altruistic strategy across the entire range of values of *h* (Fig.5b). The other dominant transitions (SC→UD→UC→SC) occur in cycles across the entire range of value of *h*, unlike the static network case where these dominant transitions were restricted to the *h* < *h*_*c*_ regime. An explanation for such category transition dynamics can be found in the manner in which rewiring affects the average degree of cooperators and defectors and the total number of C-C, C-D and D-D links. The network restructuring mechanism is such that cooperators are more likely to form links with other cooperators and break links with defector partners. Such restructuring of social ties induces frequent changes in the average degree of both altruistic and selfish agents. In the short time-scale, a rapid increase in cooperator fraction is strongly correlated with increase in the number of C-C links as well as the average degree of cooperators and decrease in the number of D-D links and average degree of defectors (Fig.6). Short time-scale fluctuations in all of these quantities are observed across almost the entire range of values of *h* but large values of cooperator fraction and number of C-C links are sustained for longer periods (as is evident from Fig.6). This allows cooperators to increase their payoffs by leveraging the larger number of connections with other cooperators in the neighborhood, whenever possible, thereby ensuring the dominance of cooperators after long-time averaging.

**FIG. 6.**
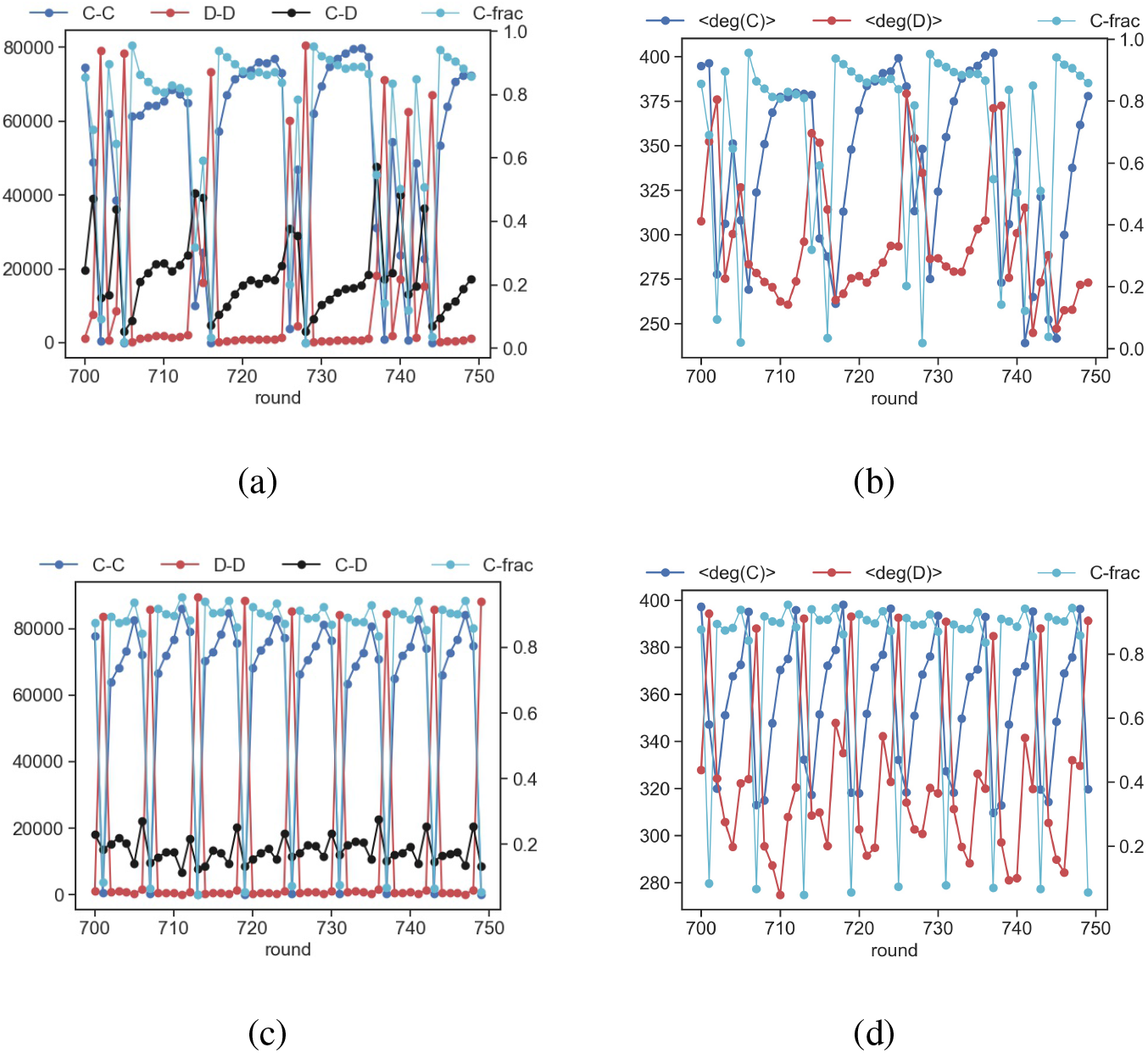
Short time-scale evolution of cooperator fraction (C-frac), the number of C-C, C-D and D-D links and the average degree of cooperators and defectors for a population with (a,b) *h* = 0.4 and (c,d) *h* = 0.9 with *ρ* = 0.3. The other parameter values are *N* = 500, *β* = 1, *ε* = 0.02, *r* = 2, *p*_*r*_ = 0.87, *p*_*b*_ = 0.7, *p*_*m*_ = 0.93, *p*_*e*_ = 0.3 and *p*_*s*_ = 0.2. C-frac denotes the fraction of cooperators.

Intriguingly, for very high habituation values (*h >* 0.7), cooperators dominate for an optimal range of values of the network rewiring probability. For *static* networks and networks with very high rewiring probabilities (bottom right and top right regions of the heat maps in Fig.5), the dominance of defectors in the population is correlated with higher satisfaction levels for defectors as seen in Fig.5c. Rewiring of all the links in every round ensures that the category transition cycle (SD→UC→UD→SD) dominates all other category transitions for large values of habituation factor (See Fig.7). Even though the network rewiring rules encourage formation of C-C links and breaking of C-D and D-D links, allowing a large fraction of nodes to restructure their connections in every round increases the possibility of formation of many C-D and D-D links. In fact, for high habituation an increase in the number of C-C links is also correlated with the increase in number of C-D links and average degree of both cooperators and defectors (see Fig. 8a,b) that makes cooperators amenable to exploitation by selfish neighbours. This leads to a fall in their payoffs below their aspiration levels, which for high *h* is calibrated to the latest payoff, thereby increasing their likelihood of defection in the next round and triggering an increase in the number of D-D links. Such large-scale rewiring of the network together with rapidly fluctuating aspiration of the players make the C-C links unstable (see Fig.8a) even over relatively short time-scales. This lack of persistence of C-C links as well as the increase in C-D links associated with increase in degree of cooperators prevents the latter from taking advantage of the ephemeral links with altruistic neighbors to increase their payoffs. Such cyclical dynamics explains the dominance of defectors in the high *h*, high *ρ* domain.

**FIG. 7.**
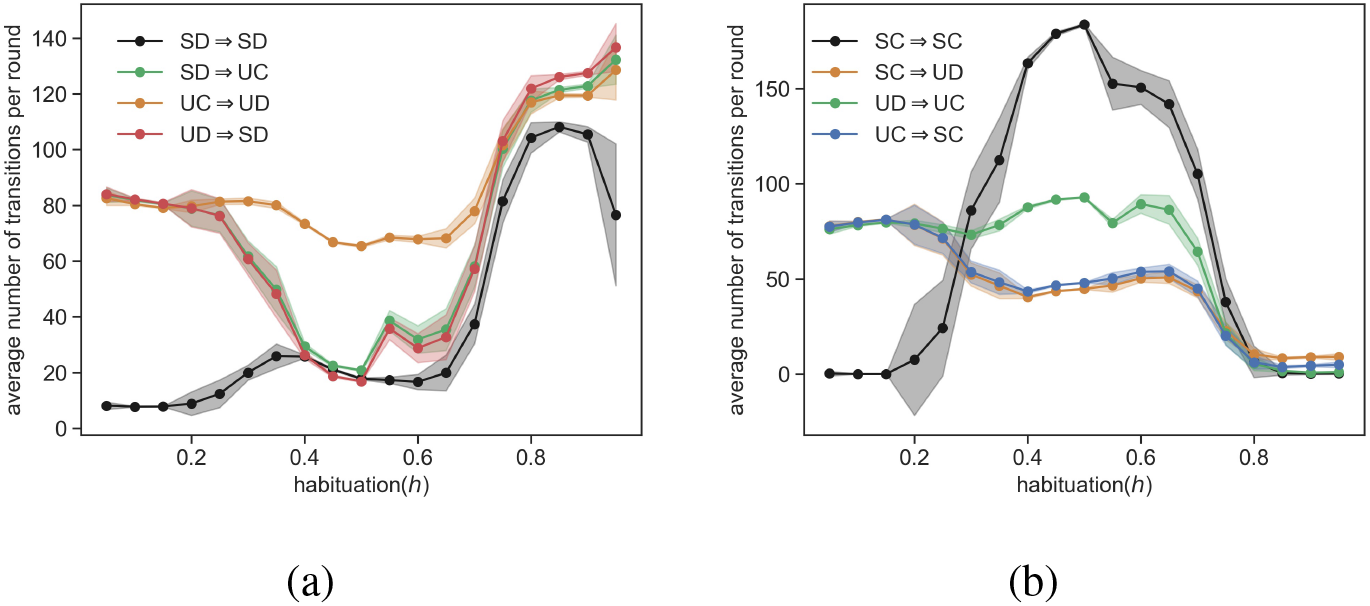
The dominant transitions including the transition cycles in the high *ρ* regime that start from (a) the SD category and (b) SC category; shown for *ρ* = 1. Each data point is obtained by first time averaging from round 500 to 750 for each trial and then averaging over 10 trials. The shaded region denotes one sigma deviation from the mean. The other parameter values are *N* = 500, *β* = 1, *ε* = 0.02, *r* = 2, *p*_*r*_ = 0.87, *p*_*b*_ = 0.7, *p*_*m*_ = 0.93, *p*_*e*_ = 0.3 and *p*_*s*_ = 0.2.

**FIG. 8.**
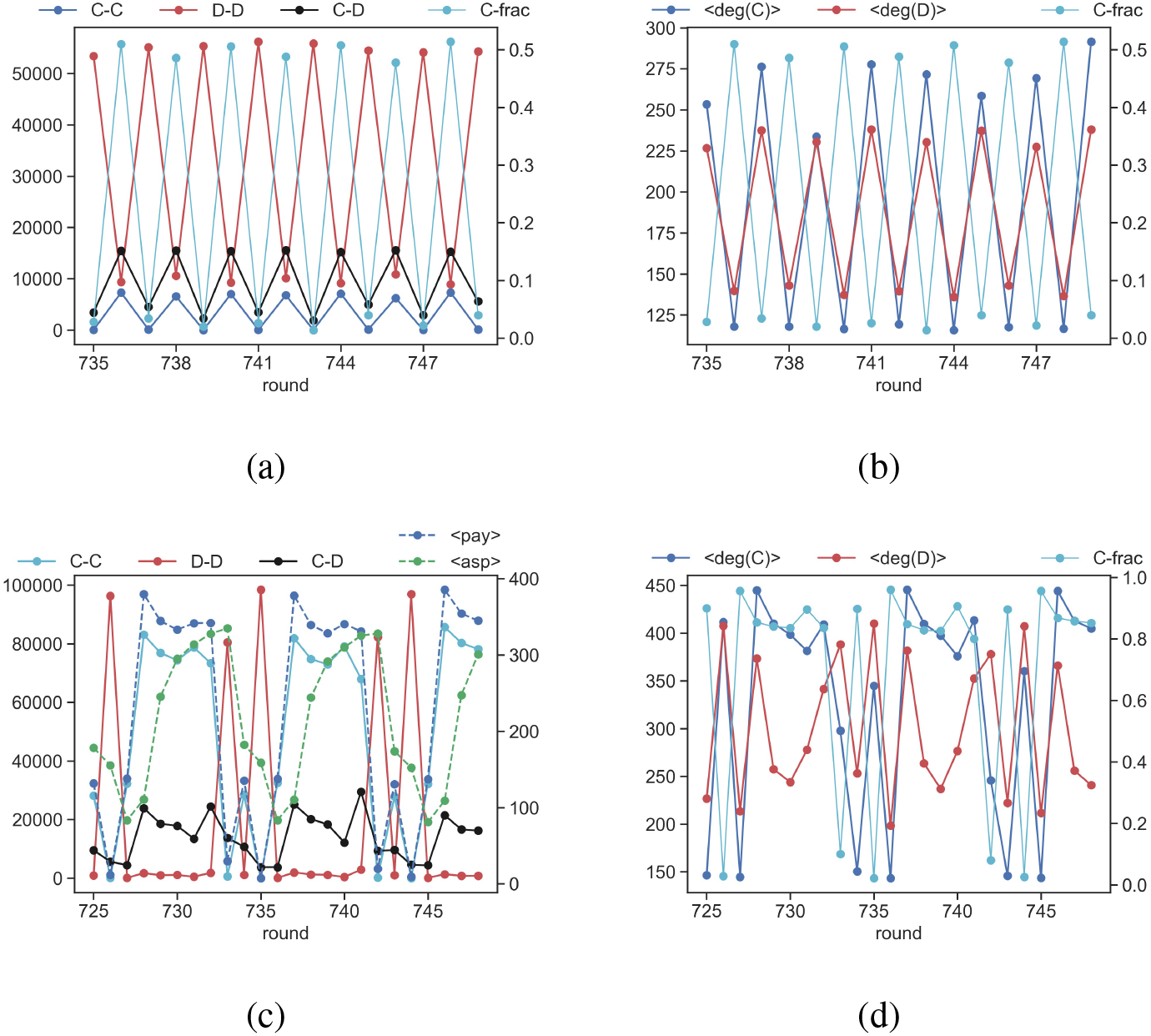
Short time-scale evolution of the number of links of different type and average degree of cooperators and defectors with *ρ* = 1.0 for (a,b) *h* = 0.9 and (c,d) *h* = 0.5. The cooperator fraction (C-frac) is shown in (a) and average aspiration and average payoff are in (c) for comparison. The other parameter values are *N* = 500, *β* = 1, *ε* = 0.02, *r* = 2, *p*_*r*_ = 0.87, *p*_*b*_ = 0.7, *p*_*m*_ = 0.93, *p*_*e*_ = 0.3 and *p*_*s*_ = 0.2. C-frac denotes the fraction of cooperators.

For a rapidly changing network characterized by very high rewiring probabilities (*ρ ≥* 0.8), there also exists an optimal range of habituation values for which cooperators dominate (see Fig. 5a). Players with a moderate habituation (top-middle region of the heat maps in Fig.5) can support cooperation at high probability of rewiring due to the large jumps in C-C links, ensuring payoffs to cooperators are on the average much higher compared to their aspirations (see Fig.8c,d). The moderate habituation values result in a slower increase in aspirations allowing cooperators to dominate for longer periods due to their larger payoffs obtained by leveraging the large number of C-C links and consequently larger average degree.

#### Polarized Aspiration

For very low habituation values, both cooperators and defectors are unsatisfied with their payoffs on the average when the network rewiring probability is very large (see Fig.5b,c) and the population is almost equally divided between cooperators and defectors (Fig.5a). In this regime, players determine their aspiration by weighing payoffs obtained in the distant past. This feature, coupled with a network that is extensively restructured in every round, spontaneously separate players into two groups based on their aspiration levels (top panel of Fig.9a). One group consists of players having aspirations peaked around 50, while the players in the other group has aspirations closer to 200. This disparity is maintained because of the inequality in the payoff received after every three rounds (see the maroon bars in the bottom panel of Fig.9a), when the players with low aspirations cooperate (in round 541) after being satisfied (in round 540) and those with high aspirations defect (in round 541) after being unsatisfied (in round 540). These actions (in round 541) leave the C’s unsatisfied and D’s satisfied with their payoffs which induces the D’s to retain their strategies and the C’s to change to a selfish strategy (in round 542). Such large-scale defection (in round 542) reduces the payoff for all players leaving everyone unsatisfied (compare light green bars in the top and bottom panel of Fig.9a) and induces everyone to change to an altruistic strategy in round 543. This restarts the cycles all over again.

**FIG. 9.**
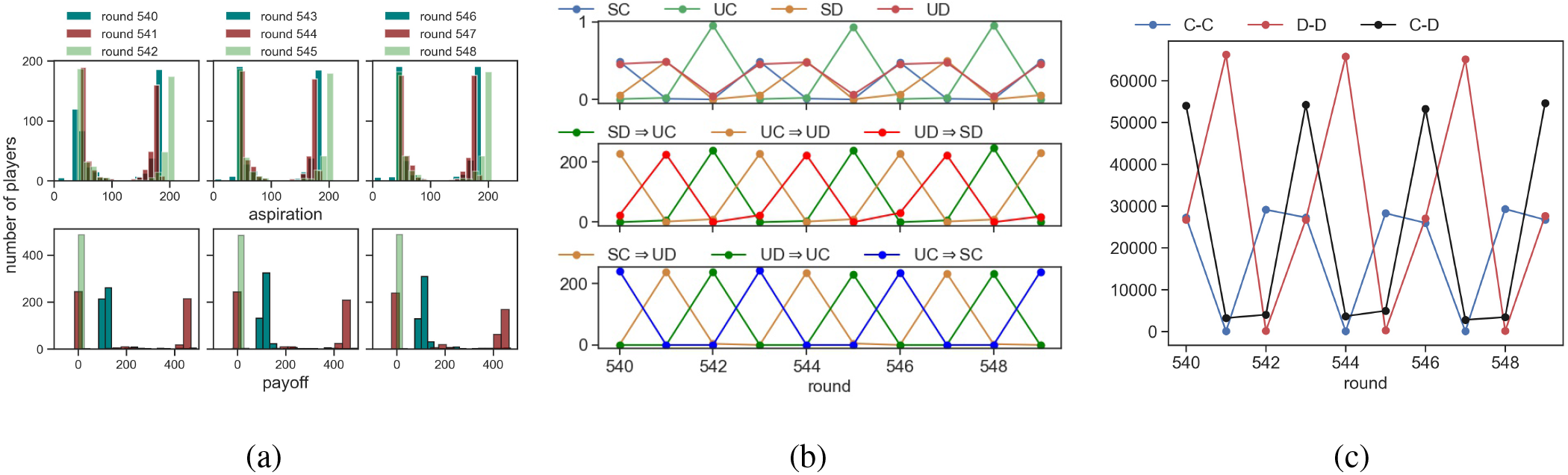
Low habituation and high restructuring probability divides the players into two distinct groups with different aspiration levels (top row) shown over 3 sets of 3 successive rounds. The histogram of payoffs obtained over the corresponding set of 3 rounds are shown in the lower row. (b) Fraction of players in each category (top row); Dominant cyclic transition between categories SD→UC→UD→SD (middle row) and SC→UD→UC→SC (bottom row) c) The short time-scale dynamics of the abundance of different types of links in the network; for *h* = 0.1 and *ρ* = 1. The other parameter values are *N* = 500, *β* = 1, *ε* = 0.02, *r* = 2, *p*_*r*_ = 0.87, *p*_*b*_ = 0.7, *p*_*m*_ = 0.93, *p*_*e*_ = 0.3 and *p*_*s*_ = 0.2.

Such a polarization in aspiration drives two sets of transition cycles (SD→UC→UD→SD and SC→UD→UC→SC) to occur simultaneously (see Fig.9b, middle and bottom panel). The SD→UC and UD→UC transitions occur in unison after defection on a large-scale reduces the payoff leaving all the players unsatisfied. These transitions increase the fraction of UC in the population to unity. Subsequently, some of these UC find that their payoffs after cooperating remain below their aspirations, prompting them to defect in the next round. The remaining fraction of UC players are satisfied with their payoffs after their altruistic act and hence retain their strategies in the next round. The increase in cooperator fraction increases the number of C-C and C-D links (Fig.9c) leaving the cooperators susceptible to exploitation by their selfish neighbors, thereby reducing their payoffs and inducing them to defect in the subsequent round.

### C. Effect of network structure

Even though the initial network topology was taken to be a random (ER) graph, we also analyzed the model by taking the initial network topology to be (i) a scale-free network and (ii) a small-world network (Fig.S1(a) and Fig.S1(b) in the supplementary material). Simulations on the above networks without rewiring produced results which are similar to Fig1. Hence, changing the initial network structure does not significantly affect our results. This is consistent with the results of a comprehensive analysis by Suri and Watts^5^ on the effect of changing network topology on cooperation levels. Their analysis, based on the results of behavioral experiments using an online platform, considered various network topologies such as cliques, small-world, random-regular networks etc. They found that the average contributions per round of players playing a PGG over many rounds did not depend on the network topologies the players were embedded in^5^. As explained in the previous paragraphs, the short time-scale fluctuations in cooperation fraction can be understood in terms of the changes in the number of C-C, C-D and D-D links as well as the degree of cooperators and defectors in the network. Nevertheless, it is interesting to ask how the specified rules of network restructuring (see Section II.B) affects the underlying network structure as characterized by evolving network metrics.

Fig.S2 in the supplementary material shows how the degree distribution of the network changes in the short time-scale as the population evolves from the defector-dominated phase (at round=717) to the cooperator-dominated phase (at round=720). In the former state the distribution is broader with the peak of the distribution occurring at a lower value of degree compared to the latter case. This skew in the degree distribution can be understood from the fact that network restructuring leads to cooperators being more connected to other cooperators in the network on an average. In an ER network, the number of C-C, D-D, C-D links is proportional to 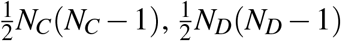 and *N*_*C*_*N*_*D*_ respectively^44^, where *N*_*C*_ and *N*_*D*_ are the number of cooperators and defectors. The restructuring of the network preserves this scaling relation between the number of C-C, D-D and C-D links and number of cooperators, defectors and product of cooperators and defectors respectively (Fig.S3 in the supplementary material). This indicates that the ER nature of the network is maintained despite network restructuring. In fact, such scaling behavior is observed even if the initial network topology is taken to be a scale-free or a small world network (results not shown) indicating that the mode of rewiring transforms the network topology to that of an ER network over time.

### D. Dynamic network: Rewiring based on satisfaction

The tendency of satisfied players to establish more connections with other satisfied players bolster cooperative behavior for a large range of learning abilities (*h ≤* 0.9), provided players rewire their connections at a moderate rate (see Fig.10). Such a rewiring mechanism also leads to a cyclic increase and decrease in C-C, C-D and D-D links but the time scales of growth and decay differ for the different types of links. When the average aspiration exceeds the average payoff, more players are unsatisfied. Since the likelihood of two unsatisfied players forming or retaining a link is low, this leads to a sparsely connected network, with players rapidly switching their strategies from the unsatisfied cooperator to unsatisfied defector (Fig.11a). Such dynamics results in a steady drop in the aspiration levels (Fig.11b) until the payoffs exceed the aspiration making players in the population satisfied with their payoffs, thereby encouraging new link formation with other satisfied players.

**FIG. 10.**
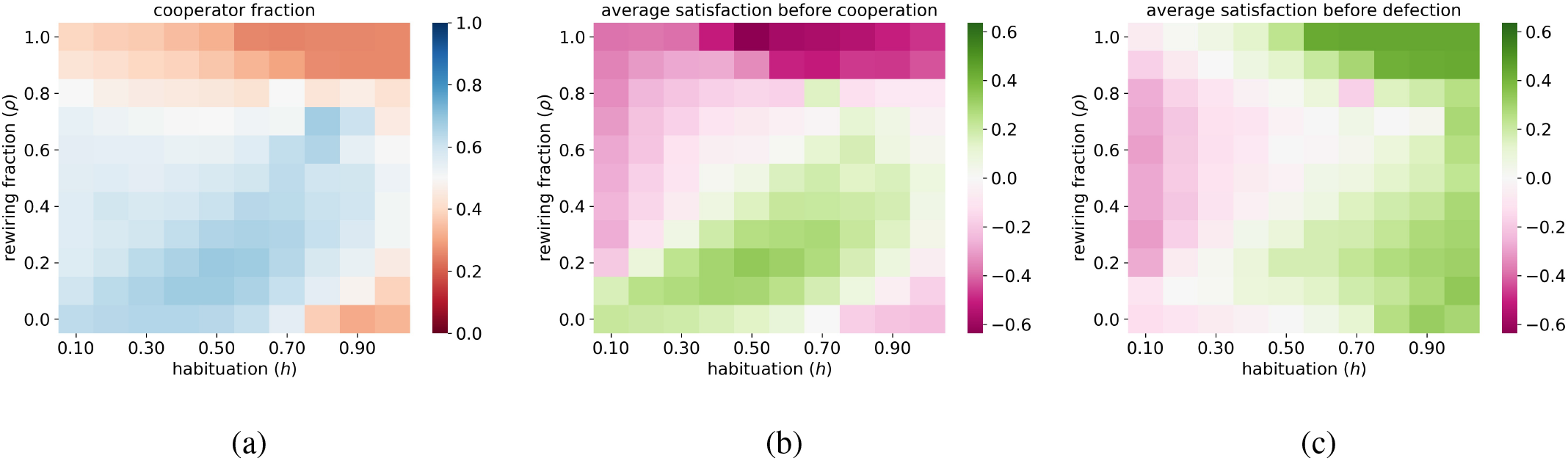
Rewiring based on satisfaction: Heat map showing the (a) cooperator fraction; average satisfaction of all individuals who (b) cooperate and (c) defect in the subsequent round, as a function of habituation (*h*) and rewiring probability (*ρ*). The other parameter values are *N* = 500, *β* = 1, *ε* = 0.02, *r* = 2, *p*_0_ = 0.2 and *p*_1_ = 0.8.

**FIG. 11.**
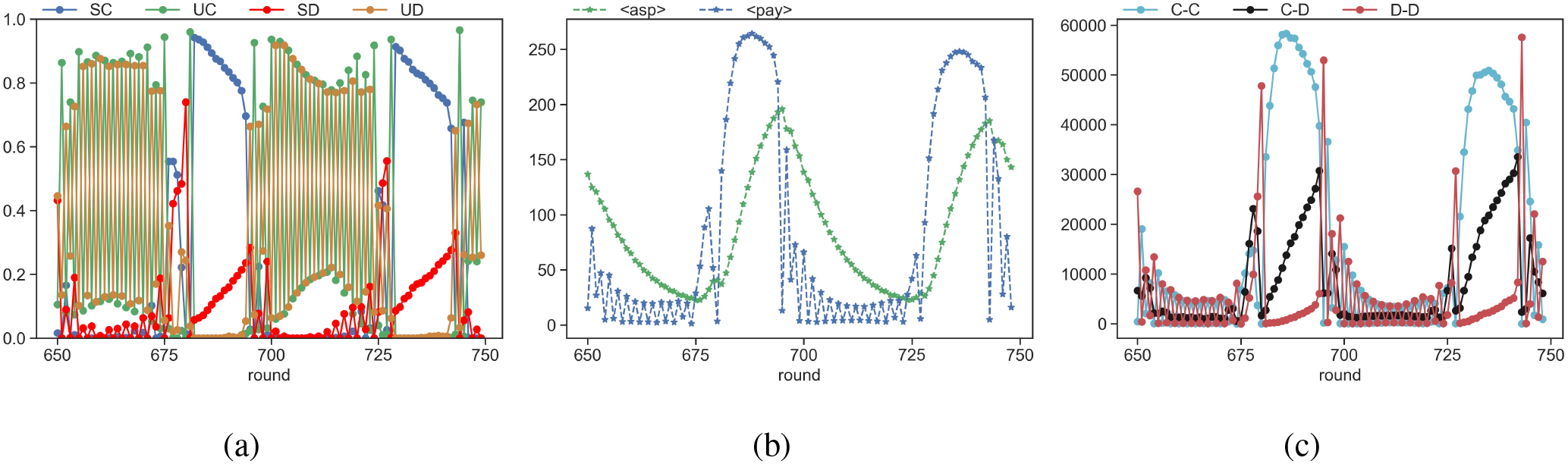
The temporal evolution of (a) the fraction of players belonging to different categories, (b) average payoff and aspiration and (c) the number of C-C, C-D and D-D links at *h* = 0.1. The other parameter values are *N* = 500, *β* = 1, *ε* = 0.02, *r* = 2, *ρ* = 0.3, *p*_0_ = 0.2 and *p*_1_ = 0.8.

Since satisfied players can be both cooperators and defectors, such link formation leads to increase in the number of C-C as well as C-D links. But, the C-C links grow faster than the C-D links and their number remains at a larger value over longer periods of time than either C-D or D-D links (Fig.11c). Cooperators are able to take advantage of more and longer-lasting C-C links to increase their payoffs. However, the gradual increase in the number of C-D links allow some defectors to increase their payoffs by exploiting cooperator neighbors. This leads to a gradual decrease in the average payoffs of cooperators and when the average aspiration level exceeds the average payoff, the cycle is repeated. The increase in the number of D-D links leads to lower payoffs for defectors and is therefore also detrimental to the satisfaction of defectors. The chances of an unsatisfied defector forming or retaining links is quite low. This results in the number of D-D links decaying very quickly after reaching a peak (see Fig.11c). All these factors act together to ensure the dominance of cooperators over larger time-scales.

For very high rewiring probabilities, cooperators have fewer links with other cooperators over long time-scales and greater likelihood of forming connections with defectors, and therefore are mostly unsatisfied. There, defectors dominate by increasing their payoffs through links with other cooperators. This explains the dominance of defectors for high rewiring probabilities for the entire range of habituation values (see Fig.10a).

## IV. SUMMARY

A large number of models of evolution of cooperation have focused on the mechanisms of direct and indirect reciprocity that requires knowledge of the actions or reputations of other members of the population. Yet, such information is often unavailable in many social conflict scenarios like PGG where individuals do not know which of their neighbors contributed to the public good and which did not. Even if individuals are aware of this information, they may still choose to change their behavior based on an internal metric. We find that a low-information reinforcement learning model where an individual’s satisfaction is based on her own aspiration levels is very effective in maintaining high levels of cooperation in a network structured PGG for a wide range of parameter values. Although fluctuations are observed over short time-scales, cooperators can dominate the population over long time-scales if their payoffs continuously exceed their aspiration levels over longer periods. This is facilitated by relatively slower changes in individual aspiration levels. If, on the other hand, aspiration is closely calibrated to the payoff (*h* > *h*_*c*_) leading to rapid changes in individual aspiration levels, defectors dominate even though cooperators are still present in the population, albeit at low fractions.

In the previous sections, the habituation factor (*h*) was considered to be a population level attribute that does not vary between individuals. However, we also considered the case where *h* is allowed to vary, thereby differentiating between the learning capabilities of different members of the population. To account for heterogeneity in learning ability, we chose *h* for each member from two distinct distributions; one in which the likelihood of selecting an *h*-value is obtained from a unimodal *β*-distribution that is peaked around *h* = 0.5 and another distribution in which the likelihood is chosen from a bimodal *β*-distribution that is peaked around *h* = 0 and *h* = 1.0. The latter case represents a situation where most members of the population have either short-term memories with aspirations calibrated to their latest payoff or aspiration levels fixed to the initial value. In both scenarios, despite short-term fluctuations, the time-averaged (over a sliding window of fixed size) fractions of cooperators exceeded defectors by about 10 − 15%; indicating that our results are robust to differences in individual learning abilities.

The effectiveness of such a model in ensuring proliferation of cooperators is enhanced if a fraction of nodes is allowed to restructure their connections by preferentially forming new links and retaining existing ones links with cooperators while breaking existing links with defectors. Such restructuring of the network is possible only if individuals possess knowledge about strategies of other players in the population. Rewiring is beneficial even in low information settings, where the strategic actions of individuals are not known to other players, if a pair of satisfied individuals (regardless of their last round actions) preferentially form new links and retain existing ones. However, rewiring provides no additional benefits relative to the static network scenario if it happens indiscriminately with nodes forming new links or breaking existing links at random (results not shown).

It is encouraging to find that a simple yet pragmatic reinforcement learning model that encourages changes in behaviour triggered by changes in individual aspiration levels, allows cooperation to not only persist but also dominate. Even a low information rewiring mechanism with a modest rewiring fraction is sufficient for cooperators to dominate over a much larger range of learning efficiencies. Our work suggests that the use of a self-defined criteria for behavioural updates, as opposed to reactive strategies that rely on actions of other interacting partners, may be quite beneficial in sustaining cooperation in social dilemmas over long time-scales.

## V. DATA AVAILABILITY

The codes and data that support the findings of this study are openly available in GitHub, at https://github.com/anuanupapa/ReinforcementLearning-StatDynNet

## VI. SUPPLEMENTARY MATERIAL

See supplementary material for additional figures.

## VII. ACKNOWLEDGEMENTS

SS is supported by a MATRICS grant (MTR/2020/000446) given by SERB, India.

## Notes

### Competing Interest Statement

The authors have declared no competing interest.

### Summary of Updates

Inserted a new sub-section analyzing the effect of of rewiring on network structure. Additional supporting figures are provided in the supplementary material file.

