## supplementary file for "Network rewiring promotes cooperation in an aspirational learning model"

**Supplementary material:** Network rewiring promotes cooperation in an aspirational learning model

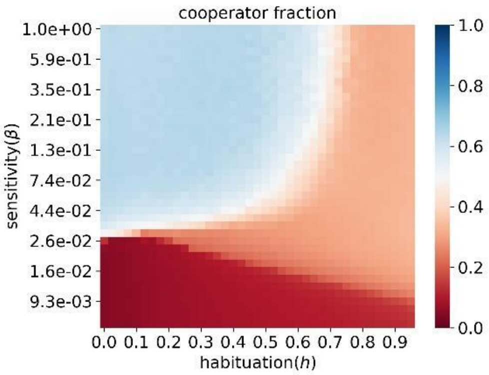


(a)

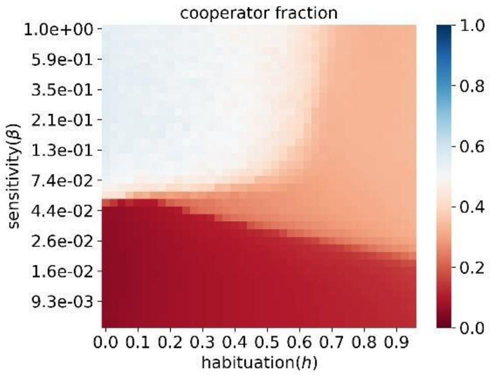


(b)

Fig. S1: Heat map showing the cooperator fraction when the initial network topology is a (a) scale-free network (b) small-world network; as a function of habituation(h) and sensitivity (β) for N = 500. Each pixel value is obtained by first averaging from round 250 to 750 for each trial and then averaging over 5 trials. The other parameter values are ε = 0.02, and r = 2.

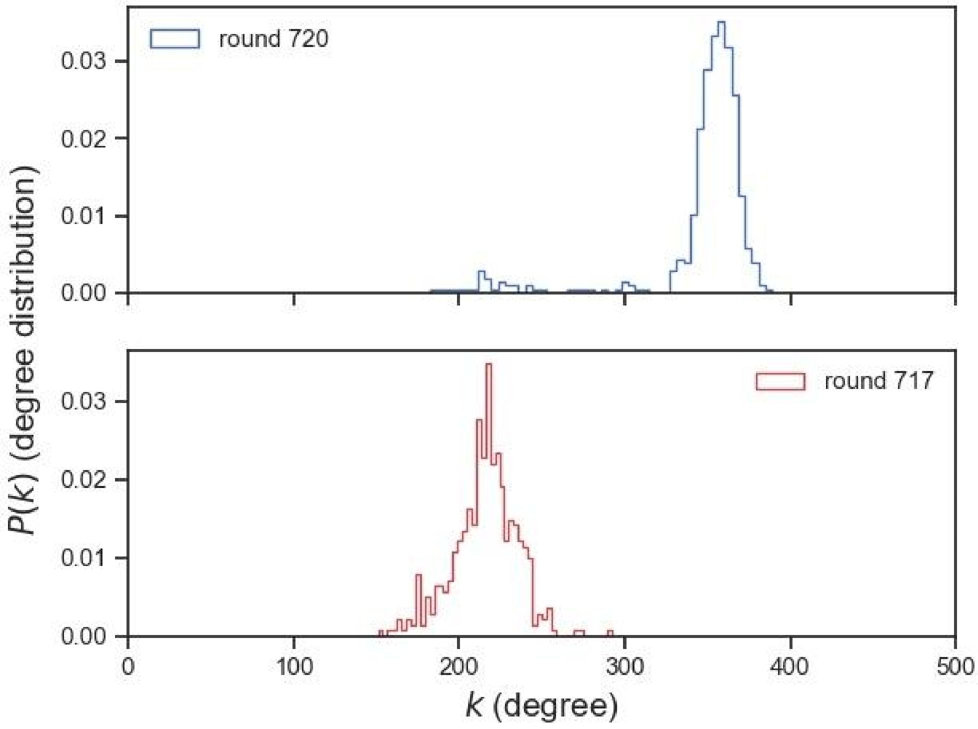


Fig. S2: The degree distribution of the network in the cooperator-dominated phase (seen in round 720) and the defector-dominated phase (seen in round 717).

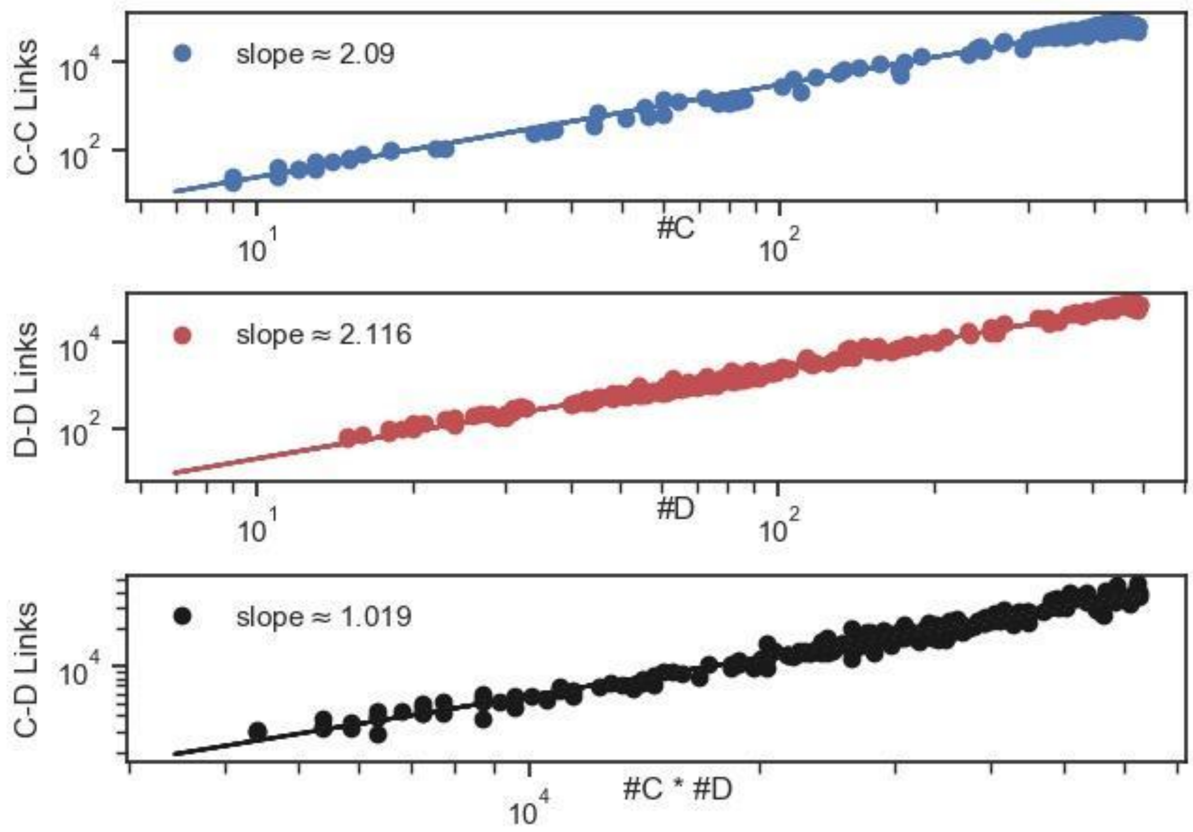


Fig. S3: The scaling of the number of (a) C-C (b) D-D (c) C-D links with the total number of cooperators, defectors and the product of C and D respectively after the system has equilibrated, for h=0.4. The data points are obtained from round 500 to 750. Other parameters are the same as in Fig.6.
